# Mating Type Gene Divergence is Associated with Life Cycle Differentiation in Scots Pine Blister Rust

**DOI:** 10.1101/2025.05.02.651906

**Authors:** Paula A. Gomez-Zapata, Christian Tellgren-Roth, Berit Samils, Jan Stenlid, Juha Kaitera, Mikael Brandström Durling, Michael F. Seidl, Åke Olson

## Abstract

Reproductive systems are central to the evolutionary and ecological processes shaping population structure and adaptability. Rust fungi are a large group of obligate plant pathogens with complex life cycles and diverse reproductive modes. In the rust fungus *Cronartium pini*, which exists in both macrocyclic heteroecious and microcyclic autoecious forms, transitions between reproductive modes can be studied within a single species. Here, we used comparative genomics to analyse the structure, diversity, and organization of mating-type (MAT) loci across both forms of *C. pini*. We identified a canonical tetrapolar system in the heteroecious form, characterized by unlinked, multiallelic homeodomain (HD) and pheromone/receptor (P/R) loci, consistent with obligate outcrossing. In contrast, the autoecious form displayed distinct reproductive signatures, including MAT gene homozygosity in some samples—indicative of clonal reproduction—and MAT gene duplications in others, suggesting self-fertility or altered mating pathways. Three HD loci were detected, but only one exhibited high allelic diversity and was consistently present, indicating functional divergence. Expanded allelic diversity at the STE3.2 pheromone receptor further suggests plasticity at the P/R locus. Although allele sharing between life cycle forms was rare, isolated cases of shared MAT alleles suggest limited historical connectivity and possible trans-specific polymorphism. Together, our findings reveal flexibility in the mating system architecture of *C. pini* and suggest that transitions between sexual and asexual reproduction may be facilitated by retained, functionally ambiguous MAT structures. This work highlights *C. pini* as a powerful model for studying life cycle evolution and mating system transitions in rust fungi.

## Introduction

Sexual reproduction dominates across eukaryotes despite its inherent costs, making its persistence a long-standing question in evolutionary biology. Although sexual reproduction imposes challenges, such as the need to find a mate and potential errors during meiosis (Otto 2009; Lehtonen et al. 2012; James 2023), it also provides significant advantages, including the elimination of deleterious mutations and the generation of genotypic diversity (Kondrashov 1994). These benefits are thought to outweigh the costs, promoting the prevalence of sexual reproduction over purely asexual reproduction (Lehtonen et al. 2012). Fungi exhibit an extraordinary diversity in reproductive modes, ranging from predominantly asexual species to almost exclusively sexual reproducers, with many species capable of alternating between both modes (Billiard et al. 2012). This makes fungi a powerful model for studying the evolution of reproduction and the genetic mechanisms underlying shifts between sexual and asexual reproduction.

Different modes of reproduction influence adaptation and long-term survival. During sexual reproduction, recombination generates new trait combinations that enhance adaptability to changing environmental conditions, including climate fluctuations and resource availability (Heitman 2015). In contrast, asexual reproduction enables rapid spore proliferation but limits genetic exchange (Taylor et al. 1999). Genetic variation in asexual fungi arises primarily through accumulation of point mutations, chromosomal rearrangements, and horizontal gene transfer (Seidl and Thomma 2014). While these mechanisms generate diversity, they are often considered less effective than recombination in maintaining standing variation that facilitates rapid adaptation. The core molecular mechanisms governing meiosis and outcrossing appear to be highly conserved across fungi, suggesting the absence of true ancient asexual lineages (Nieuwenhuis and James 2016). Consequently, asexual species have been proposed to be short-lived evolutionary “dead ends,” prone to rapid extinction (Gioti et al. 2013). However, an estimated 20% of all fungal species are thought to reproduce exclusively asexually, with no recognized sexual cycle (Hawksworth et al. 1996; Taylor et al. 2015). This paradox raises fundamental questions about how asexual fungal lineages arise and how and why they persist in nature. Understanding the genetic and ecological factors that sustain these lineages thus remains an important open question.

In fungi, sexual reproduction is controlled by mating-type (MAT) loci, which determines compatibility between individuals (Coelho et al. 2017). In fungal species with bipolar mating system, mating compatibility is controlled by a single MAT locus containing two or more idiomorphs - highly divergent alleles—that encode distinct homeodomain (HD) transcription factors involved in regulating mating and sexual development (Glass et al. 1990; Kües et al. 2011). Partner recognition is mediated by pheromone and their receptors, which in bipolar species are typically encoded outside the MAT locus. In contrast, fungal species with a tetrapolar mating system possess two unlinked MAT loci: the HD locus, which encodes transcription factors involved in post-mating development, and the P/R (pheromone/receptor) locus, which encodes pheromone precursor peptides and G protein-coupled receptors that facilitate pre-mating recognition (Kües et al. 2011). These two loci segregate independently, increasing the number of potential mating types. The HD locus is typically multiallelic, whereas the P/R locus has been reported as either biallelic or multiallelic across fungal lineages (Kües et al. 2011). This expanded mating-type architecture promotes outcrossing, reduces inbreeding, and contributes to greater genetic diversity within fungal populations. As such, elucidating the structure and allelic diversity of MAT loci is essential to understanding how reproductive systems evolve and are maintained across diverse fungal lineages.

MAT loci in rust fungi (Pucciniales, Basidiomycota) provides a valuable system for studying mating system evolution, as rust fungi exhibit an exceptional diversity of reproductive strategies (Aime et al. 2017). Rust fungi are a species-rich group of obligate biotrophs, relying on living plant tissue for survival and reproduction. Many species require multiple host species to complete their sexual life cycle, adding further complexity to their reproduction. The life cycle of macrocyclic rust fungi involves five spore stages: spermatia (0), aeciospores (I), urediniospores (II), teliospores (III), and basidiospores (IV). Macrocyclic, heteroecious rust species alternate between two taxonomically unrelated host plants: an aecial host, where plasmogamy occurs and dikaryotic spores are produced, and a telial host, where both asexual reproduction and sexual reproduction (karyogamy and meiosis) take place. Plasmogamy occurs at the aecial stage, where two haploid (n) monokaryotic individuals (spermatia) undergo fusion, forming a dikaryotic (n+n) mycelium. This mycelium produces dikaryotic aeciospores, which infect the telial host where clonal propagation occurs through urediniospore production, which re-infects the telial host. Under appropriate environmental conditions teliospores are formed, where karyogamy and meiosis occur, ultimately producing haploid basidiospores that restart the cycle. While macrocyclic, heteroecious rusts retain the full five-stage life cycle, many species exhibit life cycle reductions, including demicyclic, hemicyclic, and microcyclic forms (Jackson 1931). Additionally, some species complete their cycles on a single host (autoecious), while others remain host-alternating (heteroecious). Evolutionarily, the macrocyclic heteroecious life cycle is thought to be ancestral (Jackson 1931), with shortened life cycles representing derived states, a pattern formalized as Tranzschel’s Law (Cummins and Hirastuka 2003; Scholler et al. 2019). However, the genetic and ecological mechanisms driving these transitions remain poorly understood, leaving fundamental questions unanswered in rust fungal evolution.

The rust genera *Cronartium* and *Peridermium* represent a morphologically homogeneous group of rust fungi that infect stems, branches, and cones of *Pinus* species across North and Central America, Asia, and Europe (Samils and Stenlid 2022). Several species within these genera are important forest pathogens, causing severe economic damage to pine forests throughout the northern hemisphere (Wulff et al. 2012). Historically, *Cronartium* and *Peridermium* species have been classified separately based on life cycle differences. *Peridermium* species have been recognized as autoecious rusts, completing their entire life cycle on a single host, while *Cronartium* species have been identified as heteroecious rusts, requiring two distinct hosts to complete their life cycle (Samils and Stenlid 2022). However, molecular phylogenetic studies have led to taxonomic revisions, synonymizing certain *Peridermium* species under *Cronartium* (Hantula et al. 2002). For instance, *Peridermium harknessii* was reclassified as *Cronartium quercuum* f. sp. *banksianae* and *Peridermium bethelii* was synonymized with *Cronartium comandrae* (Vogler and Bruns 1998). These reclassifications highlight the close evolutionary relationships between autoecious and heteroecious forms within the same species. This raises fundamental questions on the evolution of these forms, e.g., if there is ongoing exchange of genetic material between the autoecious and heteroecious forms or if distinct life cycles promote genetic divergence and speciation. Understanding the genetic basis of these forms, particularly through comparative analyses of their MAT loci, provides a unique opportunity to explore how life cycle variation and reproductive modes contribute to speciation and adaptation in rust fungi.

*Cronartium pini*, the causative agent of Scots pine blister rust, exemplifies the coexistence of asexual reproducing autoecious and sexually reproducing heteroecious forms within the *Cronartium* genus (Samils and Stenlid 2022). Scots pine blister rust is most easily recognized during aeciospore sporulation, which manifests as conspicuous orange blisters on the stems or branches of infected pine trees. The heteroecious form (syn. *Cronartium flaccidum*) alternates between pine and herbaceous host plants, following a macrocyclic life cycle that includes five spore stages, including sexual reproduction via telial and basidial stages. In contrast, the autoecious form (syn. *Peridermium pini*) completes its entire life cycle on pine alone, producing only spermatia and aeciospores without known sexual reproduction. Despite these ecological and reproductive differences, both forms produce morphologically identical aeciospores with two nuclei (n+n), making them indistinguishable without molecular markers (Samils et al. 2011). Microsatellite-based studies have identified differences in heterozygosity levels between the two forms, suggesting underlying genetic divergence (Samils et al. 2011; Samils et al. 2021). However, these markers do not provide insight into the genetic mechanisms shaping these differences. Investigating the MAT loci of *C. pini* provides a unique opportunity to explore the genetic basis of its distinct reproductive strategies.

By studying the MAT loci of the two forms of *C. pini*, here we aim to address two key questions: (1) How is MAT locus diversity shaped by the reproductive strategy of each form? and (2) Are there distinct evolutionary signatures associated with each life cycle form? We predict that the heteroecious form, which undergoes sexual reproduction, will exhibit greater allelic diversity, whereas the autoecious form will show reduced diversity, consistent with predominant asexual reproduction. To test these predictions, we sequenced, assembled, and annotated the first dikaryotic reference genome of *C. pini*, providing a foundational resource for population genomic analyses, and characterized the mating compatibility system, identifying key differences in MAT gene composition. We deepen our understanding of the genetic and evolutionary mechanisms underlying the coexistence of these two life cycle forms.

## Results

### *De Novo* Reference Genome Assembly and Annotation of *Cronartium pini*

To establish a reliable foundation for investigating the mating-type (MAT) loci of *C. pini*, we sequenced and assembled the first high-quality reference genome for this species using an autoecious sample from Kallax, Sweden (sample ID: 44). The autoecious form was selected due to its presumed asexual reproductive strategy and high homozygosity, as indicated by previous microsatellite studies (Samils et al. 2011). This genetic uniformity simplifies genome assembly by reducing allelic variation between the two nuclei. The genome assembly spans 123 Mb across 80 scaffolds, with an N50 of 3.23 Mb and an N90 of 943 kb (Fig. 1A). The largest contig measures 6.62 Mb, and the genome has a GC content of 39% (Fig. 1A). Scaffolds four and five appear to represent complete chromosomes, with telomeric repeats at both ends, while 31 additional scaffolds contain telomeric repeats at one end (Fig. 1B), confirming the high contiguity and quality of the *C. pini* genome assembly. Genome completeness was assessed using BUSCO, which reveals 91% of complete single-copy orthologs (Fig. 1A). Repeat content analysis identified 62% of the genome as repetitive sequences (Fig. 1B), primarily transposable elements belonging to the long-terminal repeat (LTR) family of retrotransposons. Gene annotation using MAKER3 identified 16488 protein-coding genes, integrating Augustus, SNAP and GeneMark-ES, with RNA-seq hints and homology support to improve gene predictions. Functional annotation assigned 57% of the predicted genes to biological processes and pathways using InterProScan, Gene Ontology (GO), and KEGG pathway analyses. These results collectively indicate that we assembled a robust reference for studying the organization and diversity of MAT loci.

**Fig. 1.**
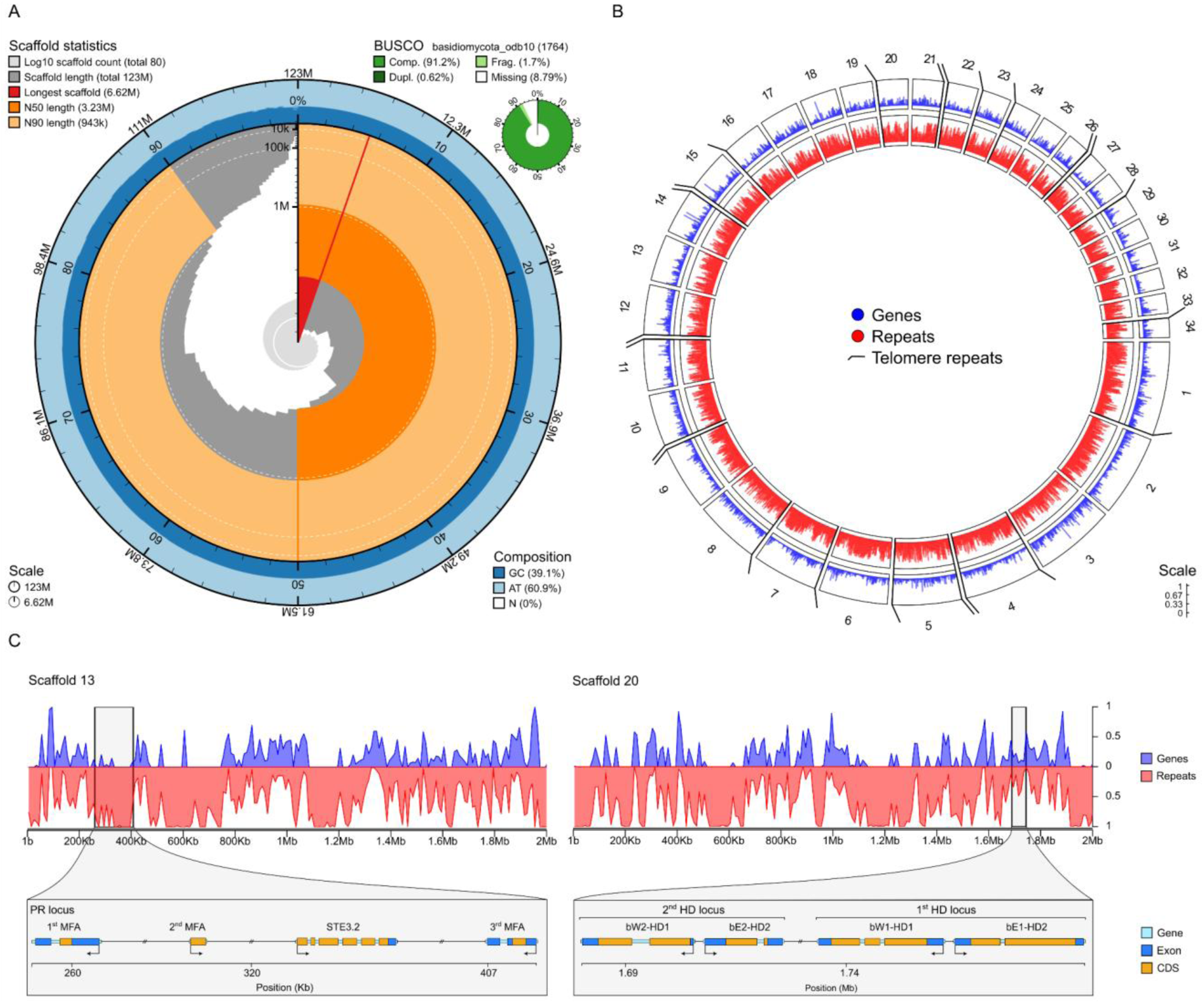
Reference Genome Assembly of the Autoecious *Cronartium pini* Life Cycle Form. **A)** Genome-wide statistics of the *C. pini* genome assembly. The circular plot illustrates scaffold length distribution. The red segment represents the longest scaffold, while all other scaffolds are arranged in size-order moving clockwise around the plot and drawn in grey starting from the outside of the central plot. Dark and light orange arcs show the N50 and N90 values. The central light grey spiral shows cumulative scaffold count. The dark vs. light blue area around the outside of the plot shows mean, maximum and minimum GC vs. AT content. BUSCO scores are shown in the upper right-hand corner. **B)** Circular representation of gene and repeat distribution (in 50 kb windows) across scaffolds that cumulatively contain 85% of the genome. Blue shows gene density, while red represents repeat elements. Thick diagonal lines mark telomeric repeats, located at scaffold ends. **C)** Mating type loci organization on scaffold 13 (P/R locus) and 20 (HD loci). The upper panel shows gene (blue) and repeat (red) density across 2Mb on each scaffold with a window size of 10 kb. The lower panel presents a detailed gene map of MAT loci, displaying the positions of pheromone receptor (STE3.2) and pheromone precursor (MFA) genes on scaffold 13, and homeodomain transcription factors (bW-HD1 and bE-HD2) on scaffold 20.

### Unlinked HD and P/R loci and Discovery of an additional HD locus in *C. pini*

Rust fungi are known to possess a tetrapolar mating system, characterized by two unlinked mating type loci: a homeodomain (HD) locus and a pheromone/receptor (P/R) locus (Cuomo et al. 2017; Luo et al. 2024). To identify the corresponding loci in *C. pini*, we searched for homologs of known MAT genes using queries from cereal rusts (Cuomo et al. 2017; Luo et al. 2024). Homologs of the P/R and HD loci are located on scaffolds 13 and 20, respectively (Fig. 1C). The physical separation of these loci onto distinct scaffolds suggests that they are located on different chromosomes. Within scaffold 20, two distinct HD loci were identified, separated by 47 kb. Each HD locus contains two transcription factor genes, bW-HD1 and bE-HD2, linked by less than 200 bp and transcribed in opposite directions (Fig. 1C). We designated the genes at the first HD locus as bW1-HD1 and bE1-HD2, and those at the second locus as bW2-HD1 and bE2-HD2 (Fig. 1C). Genome annotation and BLASTP analyses confirmed that bW-HD1 and bE-HD2 from both HD loci encode homeodomain-containing proteins, homologs to transcription factors in the Homeodomain superfamily. The P/R locus, located on scaffold 13, contains a pheromone receptor gene (PRA) of the STE3 family and three associated pheromone precursor genes (MFA) (Fig. 1C). Two MFA genes are transcribed in the reverse direction relative to the STE3 gene, located 61 kb upstream and 85 kb downstream, while the third MFA gene is transcribed in the same direction as STE3, located 14 kb upstream (Fig. 1C). The first two MFA genes exhibit high sequence similarity (94%), whereas the third MFA gene is more divergent (33% similarity). The unlinked organization of HD and P/R loci in *C. pini* aligns with the general structure of MAT loci in other rust fungi (Cuomo et al. 2017; Ferrarezi et al. 2022; Luo et al. 2024), while also exhibiting unique features such as the presence of two HD loci.

An additional STE3 gene, classified as STE3.2-1, was identified on scaffold 32. Although it belongs to the STE3 family, it was not located near any homologs of the pheromone peptide precursor genes (MFA). Due to the absence of an associated pheromone gene, STE3.2-1 was not considered part of the core MAT genes involved in mating compatibility, but it was retained for further analysis in this study.

### MAT Gene Diversity in Heteroecious Life Cycle Forms Is Greater Than in Autoecious Life Cycle Forms

To assess differences in MAT loci between autoecious and heteroecious *C*. *pini* forms, we sequenced the genomes of 18 samples (ten autoecious and eight heteroecious) using Illumina short-reads and performed *de novo* genome assemblies. Each genome assembly was queried for MAT genes using sequences from both the *C. pini* reference genome and cereal rust species (Cuomo et al. 2017; Luo et al. 2024). Homologs of bW-HD1, bE-HD2, STE3, and MFA genes were identified in all samples (Table 1). Although some MAT genes were located on separate scaffolds due to assembly fragmentation, the data provided valuable insights into MAT gene composition across samples.

**Table 1.** Mating type gene homologs identified across *Cronartium pini* samples. Sample marked with an asterisk represents the reference genome.

| Form | Sample ID | Locality | bW-HD1 | bE-HD2 | STE3.2 | MFA | STE3.2-1 |
| --- | --- | --- | --- | --- | --- | --- | --- |
| Autoecious | 44* | Kallax, Sweden | 2 | 2 | 1 | 3 | 1 |
|  | 1001 | Jokela, Finland | 1 | 1 | 1 | 3 | 1 |
|  | 1005 | Jokkmokk, Sweden | 1 | 1 | 1 | 2 | 1 |
|  | 1006 | Jokkmokk, Sweden | 1 | 1 | 1 | 2 | 1 |
|  | 543 | Halland, Sweden | 1 | 1 | 1 | 2 | 1 |
|  | 417 | Halland, Sweden | 2 | 2 | 1 | 3 | 1 |
|  | 868 | Kallax, Sweden | 2 | 2 | 1 | 2 | 1 |
|  | 952 | Pudasjärvi, Finland | 2 | 1 | 2 | 4 | 1 |
|  | 958 | Pudasjärvi, Finland | 2 | 1 | 2 | 4 | 1 |
|  | 983 | Pudasjärvi, Finland | 2 | 1 | 2 | 4 | 1 |
|  | 974 | Pudasjärvi, Finland | 2 | 1 | 2 | 4 | 1 |
| Heteroecious | 427 | Gotland, Sweden | 4 | 3 | 2 | 7 | 1 |
|  | 488 | Gotland, Sweden | 2 | 2 | 3 | 7 | 1 |
|  | 806 | Ätnarova, Sweden | 4 | 3 | 2 | 4 | 1 |
|  | 777 | Ätnarova, Sweden | 2 | 2 | 3 | 4 | 1 |
|  | 782 | Ätnarova, Sweden | 3 | 2 | 3 | 4 | 1 |
|  | 909 | Kolari, Finland | 3 | 2 | 3 | 4 | 1 |
|  | 924 | Kolari, Finland | 3 | 3 | 3 | 7 | 1 |
|  | 843 | Övertorneå, Sweden | 4 | 3 | 3 | 4 | 1 |

Our analysis revealed a notable diversity among MAT loci across the 18 samples, with distinct differences in the patterns between autoecious and heteroecious samples. Interestingly, additional MAT homologs were identified in four autoecious and all heteroecious samples (Table 1), suggesting lineage-specific expansions of MAT gene families. Among autoecious samples, most contain a single STE3.2 homolog, but all Pudasjärvi autoecious samples exhibited two STE3.2 homologs and an additional MFA homolog compared to the reference genome. Notably, all Pudasjärvi autoecious samples consistently harbor two STE3.2 and four MFA homologs, a pattern not observed in other autoecious samples (Table 1). While some autoecious samples display higher MAT gene diversity than the reference genome, heteroecious samples exhibit even greater variation. The majority of heteroecious samples contain three STE3.2 homologs (excluding STE3.2-1) and four MFA homologs, with some containing up to seven MFA homologs (Table 1). Additionally, one or two extra bW-HD1 and bE-HD2 homologs were identified in heteroecious samples, further supporting higher MAT gene diversity in this life cycle form.

### Phylogenetic Analyses of HD Transcription Factor Genes Reveal Extensive Allelic Variation at the First HD Locus

To investigate the evolutionary relationships and allelic diversity of *C. pini* HD loci, we conducted phylogenetic analyses using multiple sequence alignments of protein coding sequences. Given that bW-HD1 and bE-HD2 genes follow distinct evolutionary trajectories (Luo et al. 2024), independent trees were constructed for each to resolve genetic variation accurately. Our analysis revealed three distinct clades of bW-HD1 (bW1, bW2 and bW3) (Fig. 2A) while only two clades were present in bE-HD2 (bE1 and bE2) (Fig. 2B). The HD transcription factors encoded at the first locus (bW1-HD1 and bE1-HD2) were present in all samples from both life cycle forms, whereas the second locus (bW2-HD1 and bE2-HD2) was absent in some samples from both forms, and an additional third locus (bW3-HD1) was only identified in a subset of both forms (Fig. 2). Differences in HD loci diversity were evident between autoecious and heteroecious forms. In autoecious samples, the HD genes at the first HD locus (bW1-HD1 and bE1-HD2) were identical across all individuals, clustering within a single group (Fig. 2 colored in yellow). In contrast, the heteroecious samples at the first HD locus (bW1-HD1 and bE1-HD2) exhibited extensive allelic diversity, with several distinct alleles identified for each transcription factor (Fig. 2 colored in purple). No alleles were shared between autoecious and heteroecious samples at the first HD locus (bW1-HD1 and bE1-HD2). In contrast to the diversity at the first HD locus, the second HD locus (bW2-HD1 and bE2-HD2) and third locus (bW3-HD1) were highly conserved, each forming single monophyletic clades with no detectable sequence variation across samples and forms (Fig. 2).

**Fig. 2.**
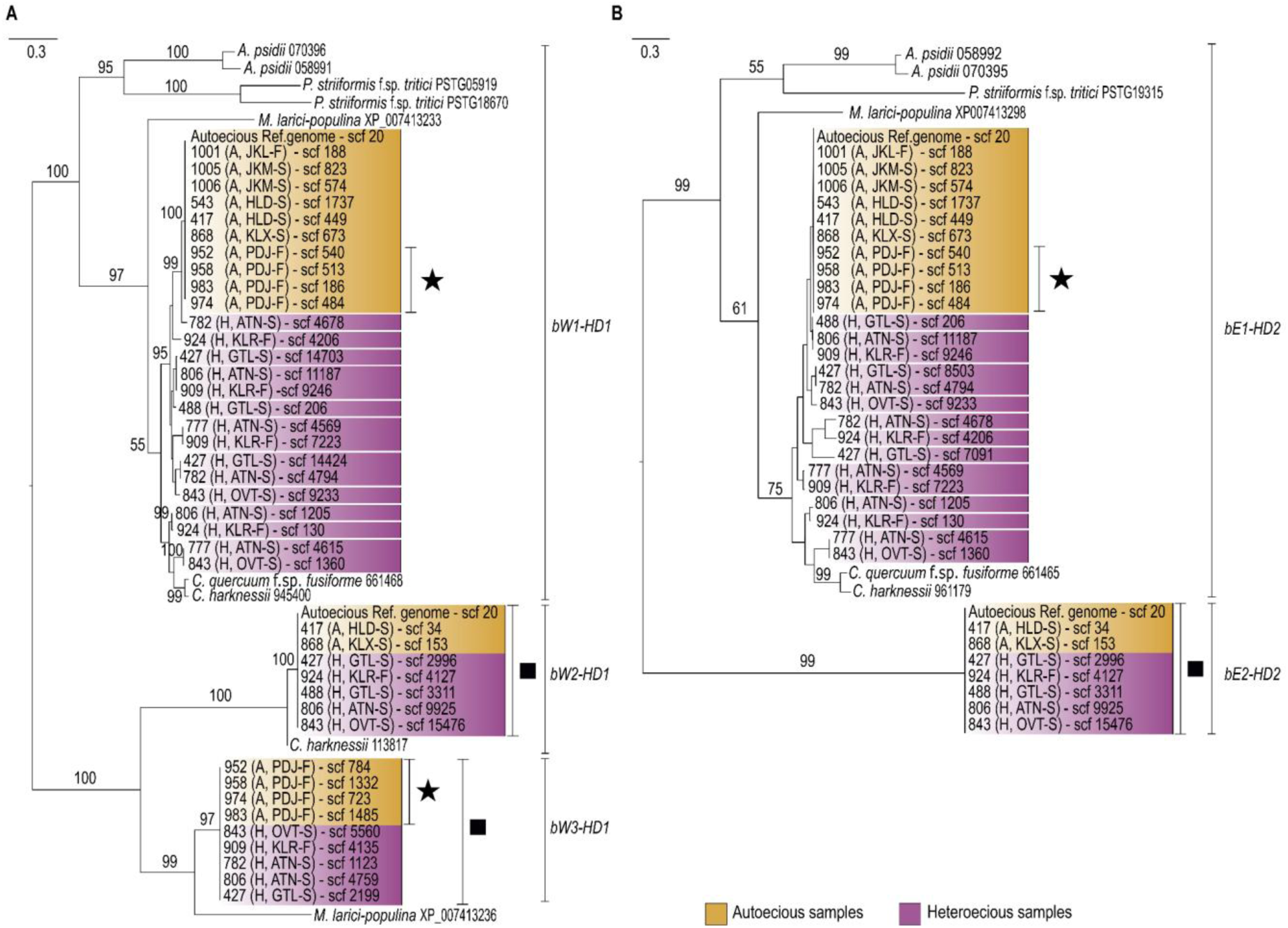
Phylogenetic Trees of bW-HD1 and bE-HD2 Protein Sequences in *C. pini* forms. Maximum likelihood phylogenetic trees from (A) bW-HD1 and (B) bE-HD2 protein sequence alignment, respectively. Ultrafast bootstrap support values are indicated at the branches. The trees are midpoint rooted. Autoecious samples are colored yellow, while heteroecious samples are purple. Sample names are formatted as follows: sample ID (A for autoecious or H for heteroecious, locality of collection [see Table 1], country of origin: S for Sweden or F for Finland), and scaffold where the HD gene was found. Black stars indicate Pudasjärvi autoecious samples and black squares indicate the clades where autoecious and heteroecious samples have identical protein sequences.

### Phylogenetic Analysis Reveals High Pheromone Receptor Diversity in *C. pini*

To investigate the allelic diversity of the pheromone receptors in *C. pini* forms, we conducted a phylogenetic analysis using the STE3.2 protein sequences of *C. pini* samples at the P/R locus. The analysis included STE3.2-1 and STE3.2 sequences from other rust species (Duplessis et al. 2011; Cuomo et al. 2017; Luo et al. 2024). Our analysis revealed high pheromone receptor diversity in *C. pini,* with twelve distinct STE3.2 groups (Fig. 3). Interestingly, no STE3.2 sequences from *C. pini* clustered with the known STE3.2-2 or STE3.2-3 sequences from *Melampsora* and *Puccinia* species (Fig. 3), demonstrating that *C*. *pini* harbors a highly divergent set of STE3.2 receptors. Sequences from the autoecious Pudasjärvi samples clustered with two known STE3.2 sequences of *C. quercuum* f. sp. *fusiforme*, forming two separate groups (Fig. 3 marked with black stars). The STE3.2-1 sequences from various rust fungi formed a strongly supported monophyletic clade, reflecting the genus-level phylogenetic relationships (Fig. 3). Within this clade, *Puccinia* sequences formed a distinct subclade, while *Melampsora* and *Cronartium* species were grouped together. We only included the STE3.2-1 sequence from the *C. pini* reference genome, since all additional STE3.2-1 sequences from autoecious and heteroecious samples were identical, suggesting that STE3.2.-1 likely does not play a role in mating.

**Fig. 3.**
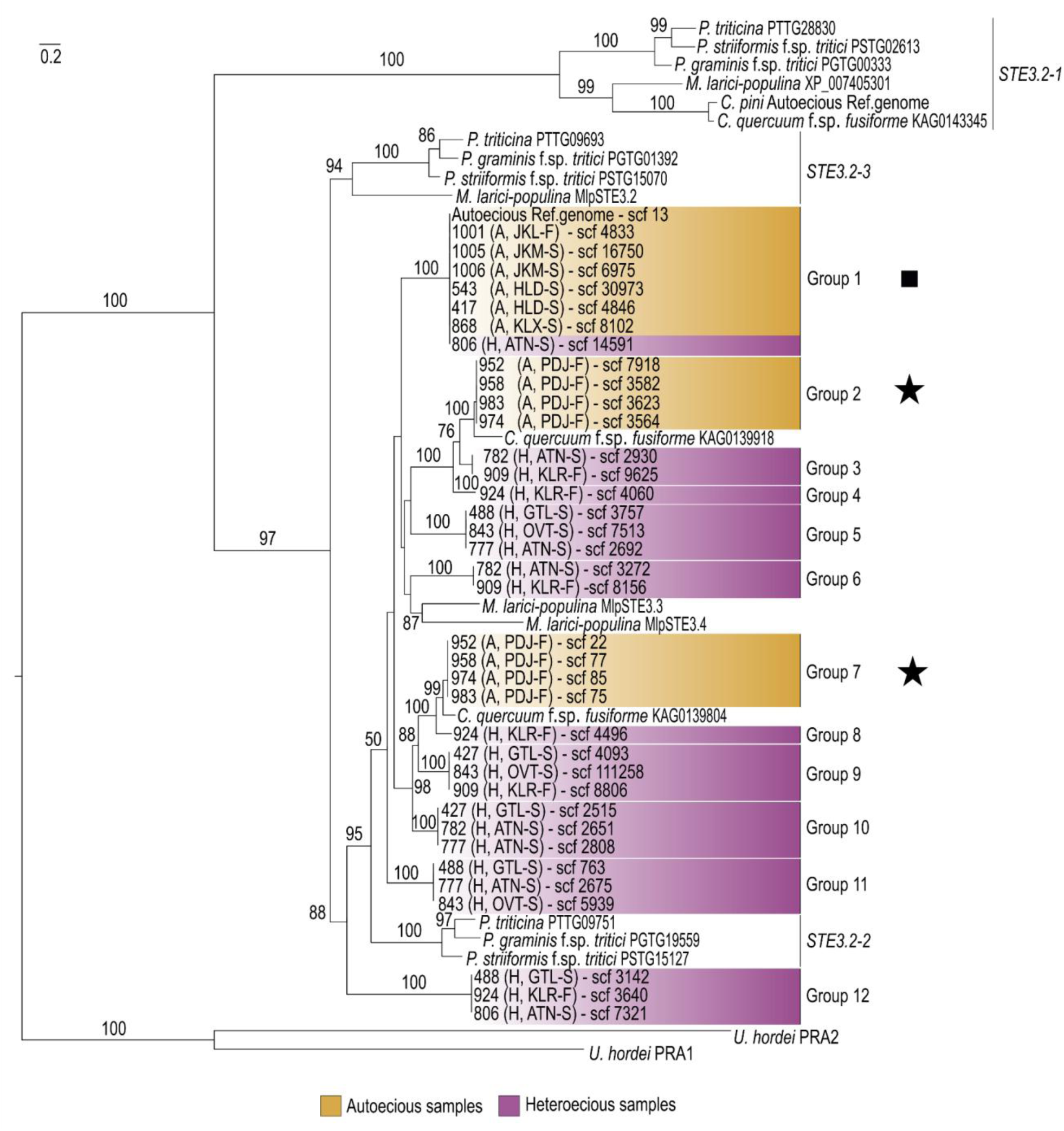
Phylogeny of STE3.2 homologs in *Cronartium pini*. Maximum likelihood phylogenetic tree from protein sequence alignments of STE3.2 genes. Ultrafast bootstrap support values are indicated at the branches. PRA sequences of *Ustilago hordei* served as outgroups. Autoecious samples are colored yellow, while heteroecious samples are purple. Sample names are formatted as follows: sample ID (A for autoecious or H for heteroecious, locality of collection [see Table 1], country of origin: S for Sweden or F for Finland), and scaffold where the STE3.2 gene was found. Black stars indicate the groups of the Pudasjärvi autoecious samples, and the black square indicates the group where autoecious and heteroecious samples have identical pheromone receptors.

Comparison of STE3.2 sequences between autoecious and heteroecious samples revealed lower diversity in autoecious samples (Fig. 3). Most autoecious STE3.2 sequences clustered within a single group (Group 1), with the exception of Pudasjärvi autoecious samples, which contained two additional STE3.2 variants grouping separately (Group 2 and Group 7, marked with black stars in Fig. 3). This distinct clustering pattern differentiates Pudasjärvi autoecious samples from the rest of the autoecious dataset. In contrast, heteroecious samples exhibited markedly higher STE3.2 diversity, with ten distinct alleles distributed across multiple phylogenetic groups. Strikingly, only one group (Group 1, marked with a black square in Fig. 3) contained pheromone receptors shared between autoecious and heteroecious samples. Within this group, autoecious samples were predominant, while heteroecious representation was limited to a single sample (806). Although this overlap was narrow, it provides a phylogenetic signal of MAT allele sharing between forms.

### Pheromone Precursors Exhibit Greater Diversity in Heteroecious than Autoecious *C. Pini* Life Cycle Forms

To explore the evolutionary diversity of mating pheromone precursors in *C. pini*, we performed phylogenetic analyses of MFA across autoecious and heteroecious samples. In this analysis, we identified three distinct MFA groups, with heteroecious samples exhibiting greater MFA diversity than autoecious samples (Fig. 4). While heteroecious samples contained MFA from all three groups, each sample consistently possessed at least one MFA from groups 1 and 2, with some also containing MFA from group 3 (Fig. 4). In contrast, autoecious samples were primarily restricted to group 2 (black circles in Fig. 4), with a few exceptions. Three autoecious samples (reference genome, 1001, and 417) contained MFA from both groups 1 and 2, whereas all Pudasjärvi autoecious samples clustered exclusively within groups 1 and 3 (black stars in Fig. 4), setting them apart from other autoecious samples. Within group 2, autoecious samples formed two closely related subgroups, reflecting lower overall diversity compared to heteroecious samples (black circles in Fig. 4). None of the autoecious samples contained MFA from all three groups. Notably, we identified a small number of clades within groups 1 and 2 that contained identical MFA sequences shared between life cycle forms (three clades marked with black squares in Fig. 4). However, even within these clades, heteroecious representation was minimal, with only a single heteroecious sample clustering alongside autoecious samples (Fig. 4, samples 427 and 806). Although rare, these shared MFA sequences suggest a signal of potential historical connectivity between mating gene pools of the two forms.

**Fig. 4.**
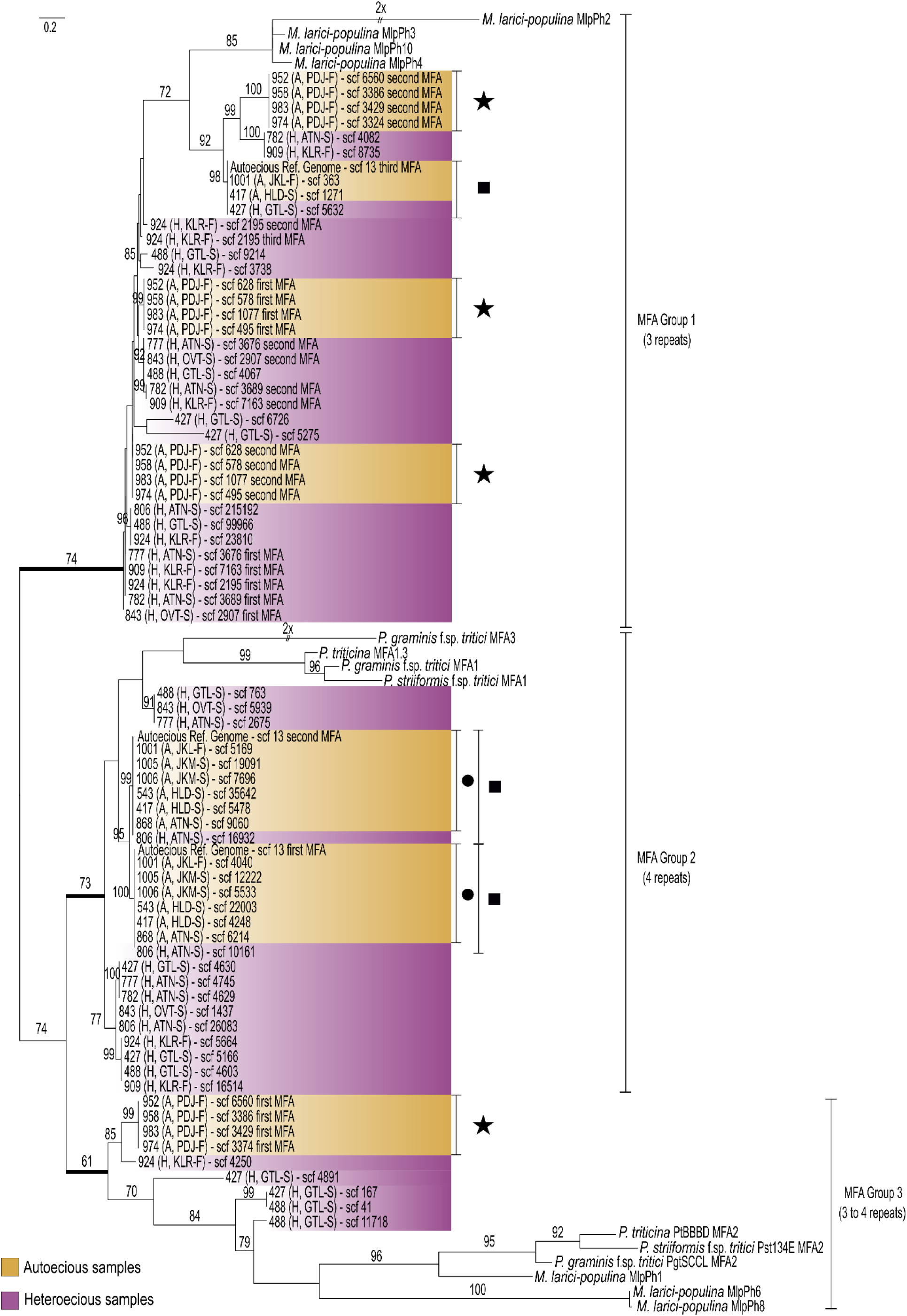
Phylogenetic Analysis of MFA Protein Sequences in *Cronartium pini* forms. Maximum likelihood phylogenetic tree from MFA protein sequence alignments. Ultrafast bootstrap support values are indicated at the branches. Thick branches denote the three MFA groups. The tree is midpoint rooted. Autoecious samples are colored yellow, while heteroecious samples are colored purple. Sample names are formatted as follows: sample ID (A for autoecious or H for heteroecious, locality of collection [see Table 1], country of origin: S for Sweden or F for Finland), and scaffold where the MFA was found. Black stars indicate the Pudasjärvi autoecious samples, black squares mark identical MFA coding sequences between forms, and black circles denote clades where most autoecious samples were clustered.

### Distinct MFA Lineages in *C. Pini* Exhibit Conserved Tandem Repeat Motifs

Tandem repeat motifs in mating pheromone precursors vary among fungal species and may influence mating efficiency and reproductive isolation (Seike and Niki 2022). To assess the repeat diversity and phylogenetic relationships of MFA in *C. pini*, we analyzed the conserved tandem repeat motifs (Supplementary Fig. S1). Each MFA lineage displayed a distinct tandem repeat structure. Group 1 has three tandem repeat units with the motifs “GNG” or “GSG,” while group 2 has four tandem repeats with the conserved motifs “GNGSHM” or “GNGSHI/GNGSWM”. In contrast, group 3 has three to four repeat units with motifs “GGGSH” or “GGSSNH” (Supplementary Fig. S1). Notably, these motifs resemble the ones found in other rust species (Duplessis et al. 2011; Luo et al. 2024), but differ in repeat number. These results suggest that while core pheromone motifs are conserved, likely due to functional constraints, repeat copy number evolves more flexibly.

### Depth of coverage patterns reveal homozygosity in autoecious and heterozygosity in heteroecious MAT loci

To determine allele composition at the identified MAT genes and assess zygosity in autoecious and heteroecious forms, we analyzed read depth across scaffolds containing these MAT loci in all genome assemblies. Genome-wide coverage distributions showed a single dominant peak in all samples, corresponding to two allelic copies—consistent with expectations for dikaryotic genomes where each nucleus contributes one allele (Supplementary Fig. S2). As a reference, we first examined the conserved STE3.2-1 gene, which consistently mapped to a single scaffold in all samples with a coverage ratio near 1.0 (Supplementary Table S1). This indicated that STE3.2-1 was present as a single allele with two identical copies in both forms, confirming it is not a duplicated or polymorphic gene and serving as a control for zygosity inference. In the autoecious samples, the read depth across scaffolds containing MAT genes also remained close to the genome-wide average (ratio ∼1.0), indicating that each MAT gene was present as a single allele with two identical copies (Fig. 5, Supplementary Table S1), consistent with homozygosity in a dikaryotic background. In contrast, heterozygosity was evident in all heteroecious samples, where reads mapped to two distinct scaffolds for the same MAT gene. These scaffolds showed a reduced depth (∼0.5×), suggesting that each scaffold harbors a distinct allele located in a different haploid nucleus (Fig. 5, Supplementary Table S1). Some MAT genes in heteroecious samples, however, exhibited a ratio of ∼1.0, suggesting that both nuclei carried the same allele or that the gene was duplicated within a single nucleus. Additionally, a few MAT genes in both forms showed elevated coverage ratios (1.5–2.0), suggesting gene duplications within one nucleus.

**Fig. 5.**
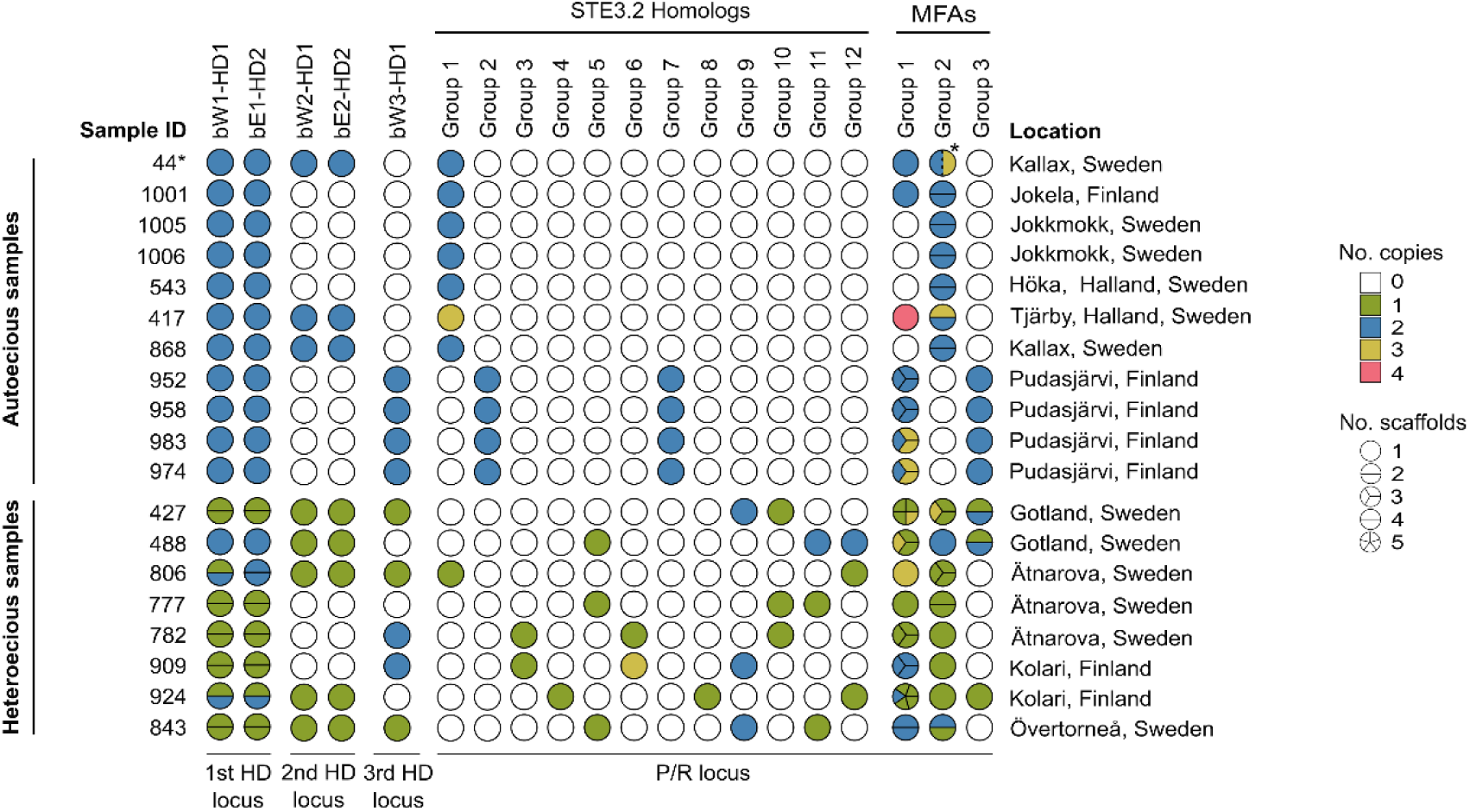
Distribution of Mating-Type (MAT) Genes across Heteroecious and Autoecious Forms of *Cronartium pini.* This figure summarizes the presence, copy number, and scaffold distribution of MAT genes across all genome assemblies. Each circle represents a specific MAT gene or gene group. The number of segments within each circle indicates the number of scaffolds on which that gene was found, while circle color denotes gene copy number, estimated from read depth (see Supplementary Table S1). Sample 44 (marked with an asterisk) represents the reference genome. In this sample, two MFA genes from Group 2 were found on the same scaffold (scaffold 13); accordingly, the group 2 circle shows two segments, despite originating from a single scaffold. On scaffold 13, the first MFA gene has two copies and the second has three copies.

## Discussion

Sexual and asexual reproduction represent contrasting evolutionary strategies, each with distinct advantages (Billiard et al. 2012; James 2023). To better understand how reproductive strategies shape genetic diversity and evolutionary trajectories, it is essential to compare the genomic signatures of these reproductive modes. Here, we used the rust fungus *Cronartium pini* as a model to investigate genetic differences associated with sexual and asexual reproduction. This fungus is ideal as it comprises two life cycle forms, a macrocyclic heteroecious form that reproduces sexually and a microcyclic autoecious form that reproduces predominantly asexually. To examine the genetic basis of these contrasting modes of reproduction, we focused on mating-type (MAT) genes, analyzing their structure, composition, and allelic diversity across both forms. Our results revealed clear distinctions in MAT gene organization and diversity between forms, reflecting divergent reproductive strategies. These findings also raise new hypotheses about the mating potential and plasticity of the autoecious life cycle to generate genetic diversity despite the apparent absence of sexual reproduction.

### Evidence for a Tetrapolar System in *C. Pini*, with Discordant Signals of Biallelism and Multiallelism at the P/R Locus

*Cronartium pini* displays characteristics of a tetrapolar mating system, with unlinked HD and P/R loci. This configuration is common across the Basidiomycota, including rust fungi such as *Austropuccinia psidii* (Ferrarezi et al. 2022), *Melampsora larici-populina* (Duplessis et al. 2011), and various *Puccinia* species (Cuomo et al. 2017; Holden et al. 2023; Luo et al. 2024). The HD locus in *C. pini* is multiallelic, consistent with the requirements of tetrapolar systems for outcrossing in rust fungi and other basidiomycetes (Kües et al. 2011; Luo et al. 2024). However, the organization of the P/R locus is more complex. While most rust fungi and many lineages of Basidiomycota (e.g., *Kwoniella heveanensis*, *Leucosporidium scottii*, *Ustilago maydis*) possess biallelic P/R loci (Kämper et al. 1995; Guerreiro et al. 2013; Maia et al. 2015), others—including *Coprinopsis cinerea* and *Schizophyllum commune*—show multiallelism at both MAT loci (Riquelme et al. 2005). In *C. pini*, the MFA gene phylogeny reveals two main compatibility groups (Fig. 4 Groups 1 and 2), suggesting a biallelic P/R structure. In contrast, the STE3.2 receptor phylogeny shows extensive allelic diversity (Fig. 3) and no clear correspondence to MFA groupings. This discordance may indicate that only a subset of STE3.2 genes participates in mating, while others are non-functional or serve alternative roles. Alternatively, *C. pini* may have a genuinely multiallelic P/R locus with decoupled evolution of pheromones and receptors. Similar patterns have been also observed in *Melampsora larici*-*populina* and *Puccinia graminis* f. sp. *tritici*, where high STE3.2 diversity complicates functional interpretation (Duplessis et al. 2011; Luo et al. 2024). While a biallelic P/R structure is considered ancestral in Basidiomycota (Maia et al. 2015), the STE3.2 diversity in *C. pini* may reflect gene duplication and divergence, as proposed for other fungi with expanded P/R loci (Riquelme et al. 2005). Functional validation—such as receptor-ligand binding studies or gene knockouts—will be essential in the future to clarify which STE3.2 genes mediate mating.

### Three HD loci in *C. pini* Suggest a More Complex Mating Compatibility Architecture

We identified three homeodomain (HD) loci in the *C. pini* genome assemblies (Fig. 2). The first locus (bW1-HD1 / bE1-HD2) is the most consistent across samples and displays high allelic diversity, making it the strongest candidate for mediating mating compatibility. In contrast, the second (bW2-HD1 / bE2-HD2) and third (bW3-HD1) HD loci were absent in some samples and exhibited no detectable sequence variation. Notably, the third locus lacks a paired HD2 gene, deviating from the canonical HD locus structure typically required for functional heterodimer formation. However, similar unpaired or rearranged HD loci have been reported in other basidiomycetes, where one HD gene exists without its corresponding partner (James et al. 2013). These non-standard architectures suggest that partial or degenerated HD loci may be evolutionarily maintained under relaxed selection or may serve as regulatory elements. Although these patterns initially suggested that only the first HD locus contributes to mating compatibility, further examination revealed that one heteroecious sample (488) lacked allelic diversity at the first HD locus but possessed genes from the second HD locus. This finding raises the possibility that additional HD loci may contribute to compatibility in certain genomic backgrounds or under specific life cycle conditions. Thus, while the first HD locus appears to be the primary determinant of mating-type specificity, the conservation of the second and third loci across life cycle forms—despite their low allelic diversity—supports the hypothesis that they may retain functional or regulatory significance. Functional validation will be essential to clarify the roles of each HD locus in compatibility and post-mating development.

### Mating Gene Profiles Reflect Distinct Reproductive Strategies Between Life Cycle Forms

The two life cycle forms of *C. pini* show contrasting mating gene profiles that reflect their reproductive strategies. Heteroecious samples exhibit a canonical tetrapolar system, characterized by high allelic diversity at HD and P/R loci. The co-occurrence of aecial and telial hosts within the same geographic regions facilitates sexual reproduction in this life cycle form. Consistent with this, read depth analysis shows that MAT genes in heteroecious samples are present at approximately half the genome-wide coverage (Supplementary Table S1), indicating that each of the two nuclei in the dikaryotic state carries a different MAT allele (Fig. 6A). Together, these patterns strongly suggest that the macrocyclic heteroecious life cycle of *C. pini* is tightly linked to a sexual, outcrossing reproductive strategy, as reflected in both mating gene diversity and nuclear composition.

**Figure 6.**
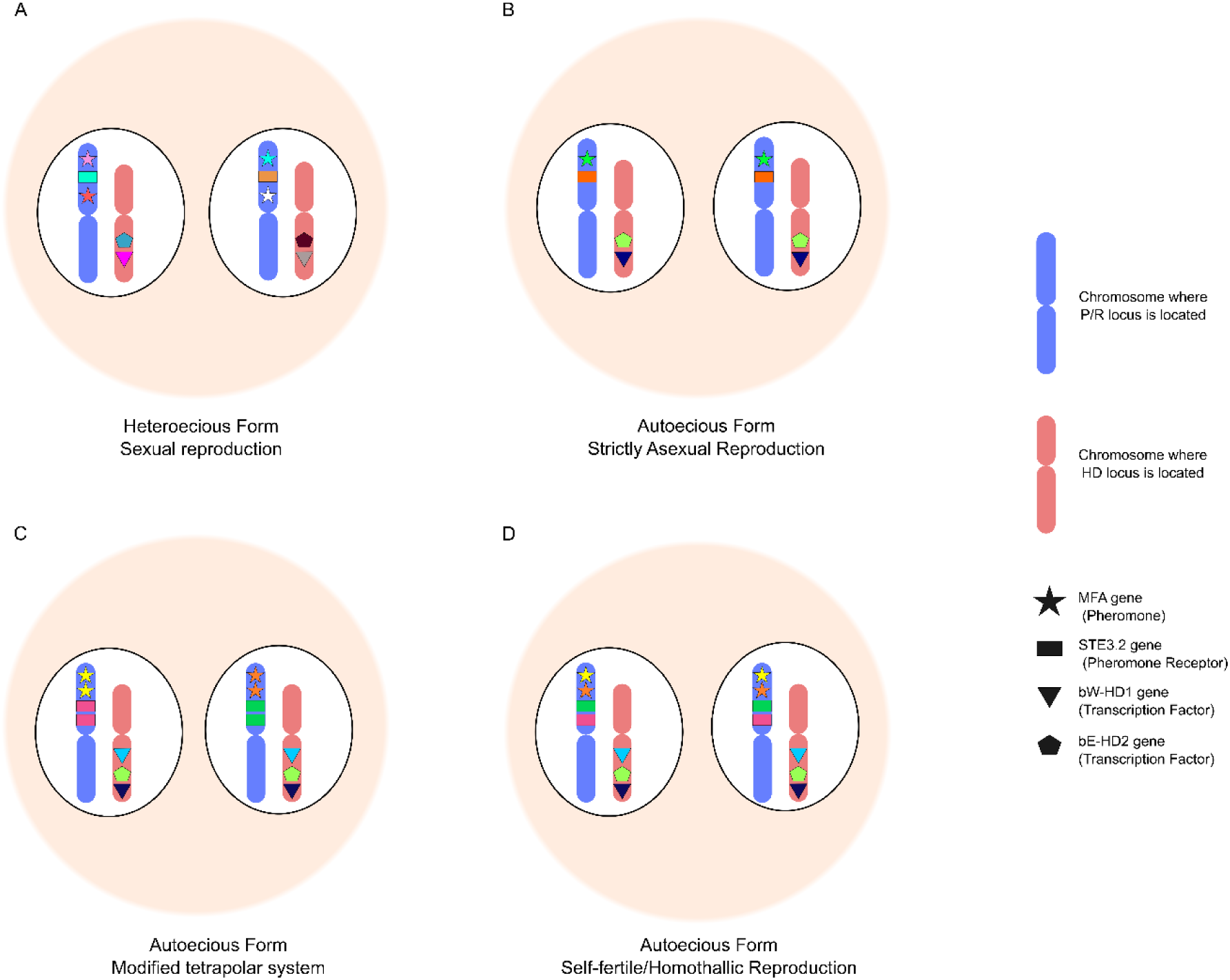
Reproductive modes inferred from MAT gene diversity in *Cronartium pini*. Schematic representation of four reproductive modes identified in *C. pini* based on MAT gene composition, allelic variation, and read-depth analysis. **A)** Sexual reproduction (Heteroecious form; typical tetrapolar mating system): Each nucleus contains distinct alleles at both MAT loci—multiallelic HD transcription factors at HD loci and variable pheromone receptor (STE3.2) and precursor (MFA) genes at the P/R locus. This configuration enables outcrossing and supports a classical tetrapolar mating system. **B)** Strictly asexual reproduction (Autoecious form; clonal propagation): Both nuclei are genetically identical at mating genes, showing no allelic variation at either HD or P/R loci. Homozygosity is observed for all MAT genes, consistent with obligate clonal reproduction and the absence of mating compatibility. **C–D)** Putative reproductive strategies in a subset of autoecious samples, from Pudasjärvi, Finland. Due to the unphased nature of the genome assemblies, the configurations represent two possible scenarios: **C)** Modified tetrapolar system (Autoecious form; gene duplication at P/R locus): Both nuclei harbor identical HD genes, but multiple copies of STE3.2 and MFA are present on each nucleus due to gene duplication. Despite HD homozygosity, the presence of different P/R genes at each nucleus may partially maintain components of the tetrapolar system, although the absence of HD allele diversity likely prevents successful post-mating development. **D)** Self-fertile (homothallic) reproduction (Autoecious form; distinct P/R alleles per nucleus): Each nucleus carries different alleles at the P/R locus (STE3.2 and MFA), while HD genes remain identical. This P/R variation within each nucleus suggests potential for self-compatibility, allowing mating within a single haploid individual, indicative of primary homothallism or facultative selfing.

In contrast, autoecious samples exhibit two distinct reproductive signatures in the mating type genes. One subset of samples (44, 1001, 1005, 1006, 543, 417, and 868) lacks allelic variation at any MAT gene, carries only a single STE3.2 allele, and has identical MAT gene alleles in both nuclei (Fig. 6B). This genotype supports a clonal reproductive strategy. Interestingly, another subset of samples (952, 958, 983, and 974) from Pudasjärvi, Finland, displays a more complex pattern. These samples contain two STE3.2 alleles, two MFA alleles (Groups 1 and 3), and two HD loci, each with two identical alleles. Given the unphased nature of these dikaryotic genome assemblies, we propose two scenarios to explain how the mating gene composition might relate to reproduction in these samples. In the first scenario, these samples represent a modified tetrapolar system with altered post-mating development (Fig. 6C). Each haploid nucleus would carry a duplicated pheromone and receptor gene distinct from each other allowing compatibility and plasmogamy. However, because all HD genes are homozygous, heterodimer transcription factors would not be able to form, unable to maintain a dikaryotic stage in post-mating development. This may lead to an unstable dikaryotic phase followed by karyogamy and rapid meiosis, mimicking teliospore and basidiospore formation. Whether the diploidization occurs in the aeciospore or during germination remains unknown. However, the resulting haploid cells produced by meiosis would inherit one of the two MAT loci configurations originally present in the dikaryotic cells.

In the second scenario, reproduction may occur via a self-fertile or homothallic mechanism (Fig. 6D). In this case, each haploid nucleus would carry two distinct pheromone and receptor genes, while the HD genes remain homozygous. The presence of compatible pheromone and receptor genes within the same nucleus would allow self-compatibility, enabling plasmogamy—consistent with primary homothallism or facultative selfing (Billiard et al. 2012). However, if only one nucleus is required, the presence of two nuclei in aeciospores remains unexplained. This may reflect an incomplete transition to homothallism or a yet-unrecognized developmental role for the second nucleus.

Our findings are consistent with recent population genetic evidence presented by Zhang and Samils (Zhang and Samils 2024), who compared heteroecious and autoecious populations of *C. pini* using multilocus genotyping. This analysis revealed high genotypic and allelic diversity in heteroecious populations—consistent with regular outcrossing—while autoecious populations showed reduced genotypic diversity and widespread homozygosity at SSR markers. These patterns support clonal propagation in autoecious forms but also suggest the possibility of self-fertilization (primary homothallism), as several genotypes remained homozygous yet genetically distinct. Zhang and Samils proposed a model in which dikaryotic aeciospores produce monokaryotic hyphae capable of selfing via fusion of compatible spermatia and receptive hyphae of identical genotype. This hypothesis aligns closely with our scenario of modified tetrapolar system in Pudasjärvi samples (Fig. 6C), in which duplicated pheromone and receptor genes within each nucleus may facilitate mating compatibility in the absence of HD heterozygosity. Together, these independent lines of evidence suggest that autoecious *C. pini* may employ other reproductive strategies than clonality, representing a potential evolutionary transition away from obligate outcrossing.

Interestingly, previous studies have reported variation in nuclear content among rust spores, including the presence of haploid and diploid stages in spore types historically classified as dikaryotic (Talhinhas et al. 2023). A similar phenomenon may apply to the autoecious *C. pini* samples from Pudasjärvi, where the dikaryotic state may not be maintained throughout development. Notably, spermatia have been observed in several Finnish populations, including Pudasjärvi (Kaitera and Nuorteva 2008), but their presence has not been investigated in Swedish populations. The occurrence of spermatia in Finland aligns with either of the two proposed reproductive scenarios and suggests that some autoecious populations may engage in reproductive strategies beyond strict clonality. Microscopic analyses of nuclear dynamics and spore germination in these populations will be essential to determine whether ploidy shifts occur during the aecial stage and to clarify the reproductive cycle of the autoecious form.

Taken together, our findings suggest that Pudasjärvi isolates may represent an intermediate stage between the two life cycle forms. The retention and duplication of MAT genes, combined with limited allele sharing and evidence for reproductive plasticity, point to a system in flux. Whether Pudasjärvi populations are remnants of an ancestral heteroecious form or represent a derived, self-compatible lineage remains an open question. However, their genetic composition provides a rare window into the mechanisms and the mating type genes role in the transitions between sexual and asexual reproduction in rust fungi.

### Potential for Mating Between Autoecious and Heteroecious Forms of *C. pini*

Our findings reveal limited allele sharing between autoecious and heteroecious *C. pini*, suggesting restricted genetic exchange between forms. However, the conservation of several MAT genes across some samples from both forms suggests that, in theory, haploid cells from each form could be compatible for mating. Notably, one heteroecious sample (806) carried identical alleles at multiple MAT genes with an autoecious sample (417), providing the strongest evidence to date of potential genetic exchange between forms. Specifically, transcription factors at second HD locus, the STE3.2 pheromone receptor from group 1, and MFA alleles from group 2 were identical between these two samples, suggesting that haploid nuclei from each form may have contributed to the dikaryotic genotype of sample 806. Alternative explanations, such as incomplete lineage sorting or ancestral polymorphism, could also explain these shared alleles without requiring direct genetic exchange. Further investigation is necessary to distinguish between these possibilities.

Interestingly, a recent population genetics study (Zhang & Samils 2024), based on multilocus genotyping, detected gene flow predominantly from autoecious to heteroecious populations. Although our study does not directly test genome-wide introgression, the shared MAT alleles observed between sample 417 and 806 are consistent with this directional signal. This raises the possibility that autoecious lineages may occasionally contribute to the genetic background of heteroecious populations, particularly in zones of past or present sympatry (Samils et al. 2021).

The geographic pattern of allele sharing between samples from different forms further complicates interpretation. Despite shared MAT alleles between sample 806 (Ätnarova, Sweden) and sample 417 (Tjärby, Halland, Sweden), no such sharing was observed in other locations where both forms occur in closer proximity. This suggests that genetic exchange, if it occurs, is rare and spatially restricted. Targeted sampling in localities where both life cycle forms co-occur would help clarify the extent and frequency of potential inter-form genetic exchange.

Overall, while our results suggest that limited allele sharing between autoecious and heteroecious *C. pini* is possible, the underlying mechanisms remain unresolved. Future studies using phased whole-genome data, including introgression and haplotype-sharing analyses, will be essential to determine whether inter-form mating occurs naturally and how it might influence population structure. Expanding geographic sampling in regions where both forms co-occur will also be critical to assess whether gene flow is ongoing or reflects ancestral connectivity.

## Conclusions

Through comparative genomics and phylogenetic reconstruction, we highlight the diversity and plasticity of mating systems within a single fungal species, *Cronartium pini*. By linking MAT gene architecture to distinct life cycle forms, we show how genomic structure reflects and potentially constraints reproductive strategies. The heteroecious form exhibits a canonical tetrapolar mating system with multi allelic diversity at both loci, consistent with obligate outcrossing and recombination. In contrast, the autoecious form suggests a spectrum of reproductive modes, from strict clonality to possible self-compatibility, and retains mating gene architectures that may facilitate occasional recombination. These findings support a dynamic view of mating system evolution, where transitions between sexual and asexual reproduction are not binary but involve intermediate states shaped by genomic architecture, ecological context, and possibly host availability. The presence of conserved but functionally ambiguous MAT gene copies, particularly in the autoecious form, raises the possibility that gene retention provides a latent capacity for mating under favorable conditions. More broadly, our work underscores how changes in mating system structure can influence evolutionary trajectories, facilitate life cycle transitions, and shape pathogen diversification. As such, understanding the genetic basis of mating in rust fungi not only informs fungal evolution but also provides a foundation for predicting long-term trajectories in pathogen adaptation.

## Materials and Methods

### Collection of *C. pini* Aeciospores

Aeciospores of *Cronartium pini* were collected from Scots pine (*Pinus sylvestris* L.) in multiple pine forests across Northern Fennoscandia during 2011–2012 (Supplementary Table S2). The collection included both autoecious and heteroecious forms, identified using SSR markers following the protocol described by Samils et al. (Samils et al. 2011). Individual aecia were sampled using tweezers sterilized with 70% ethanol and placed into separate 1.5 mL microcentrifuge tubes. To minimize cross-contamination, only closed aecia were selected. Samples were transported immediately to the laboratory and stored at –20 °C until further processing.

### DNA Extraction and Sequencing

Genomic DNA for the reference genome was extracted from at least 100 ng of aeciospores using the NucleoSpin Plant II kit (Macherey-Nagel), following the manufacturer’s protocol. Spores were suspended in PL1 extraction buffer supplemented with 10 µL RNase A and a combination of glass beads (three 3 mm and twenty 2 mm) in a screw-cap tube. Homogenization was performed using a Precellys bead beater (Bertin Technologies, Montigny-le-Bretonneux, France) for 2 × 45 s at 5500 rpm, followed by incubation at 65 °C for 10 min. DNA from 18 additional *C. pini* samples was extracted from dry-homogenized aeciospores using the CTAB method (Samils et al. 2011).

For the reference genome, a SMRT sequencing library was created from 5 µg of genomic DNA was using the SMRTbell Express Template Prep Kit 2.0 according to the manufacturer’s instructions (PacBio, Menlo Park, USA) and sequenced on a Sequel IIe instrument using the Sequel II sequencing plate 2.0, binding kit 2.2 on one Sequel® II SMRT® Cell 8M, movie time 30 hours and pre-extension time 2 hours. (PacBio, Menlo Park, USA). A total of 5 Gb of HiFi-data was obtained. For the remaining 18 samples, sequencing libraries were prepared from 50 ng of DNA using the SMARTer ThruPLEX DNA-seq Kit (Takara, cat# R400676) with Unique Dual Index Sets A–D (cat# R400665–8). DNA was fragmented to an average insert size of 350–400 bp using a Covaris E220 ultrasonicator, and library preparation followed the manufacturer’s protocol (guide #112219). Libraries were sequenced on the Illumina NovaSeq 6000 system using paired-end 150 bp reads with SP flow cells and v1.5 sequencing chemistry, including a 1% PhiX spike-in. Sequencing was performed at the SNP&SEQ Technology Platform in Uppsala, part of the National Genomics Infrastructure (NGI) Sweden and SciLifeLab. DNA quality and concentration were evaluated using FragmentAnalyzer or TapeStation and Qubit/Quant-iT, respectively. Each NovaSeq lane produced at least 650 million paired-end reads, with ≥75% of bases achieving a Phred quality score of Q30 or higher, contingent on sample QC.

### RNA Isolation and Sequencing

Total RNA was extracted from approximately 50 ng of aeciospores from the reference *C. pini* sample, stored at –80 °C. Extraction was performed using the Spectrum™ Plant Total RNA Kit (Sigma-Aldrich), following the manufacturer’s protocol. Spores were mixed with lysis buffer containing β-mercaptoethanol and a combination of glass beads (three 3 mm and twenty 2 mm) in a screw-cap tube. Homogenization was carried out in a Precellys bead beater (Bertin Technologies, Montigny-le-Bretonneux, France) for 2 × 45 s at 5500 rpm. RNA concentration was measured using a NanoDrop spectrophotometer (Thermo Fisher Scientific, Waltham, MA, USA), and residual genomic DNA was removed with DNase I (Sigma-Aldrich). RNA integrity was assessed using a Bioanalyzer DNA 7500 (Agilent Technologies, Boulder, CO, USA), and only samples with RNA integrity number (RIN) values above 5.4 were selected for sequencing. RNA-seq libraries were prepared from 500 ng of total RNA using the TruSeq Stranded mRNA Library Preparation Kit (cat# 20020595, Illumina Inc.), which includes poly(A) selection. Unique Dual Indexes (cat# 20022371, Illumina Inc.) were used for sample indexing. Library preparation was performed according to the manufacturer’s protocol (#1000000040498). Sequencing was carried out using a MiSeq System (Illumina) with v3 chemistry and paired end 150 bp read length at the Department of Medical Sciences, National Genomics Infrastructure (NGI), SNP&SEQ Technology Platform, Uppsala Biomedical Centre (BMC), Uppsala University, Sweden. The resulting transcriptome data were used for genome annotation and validation of mating-type (MAT) gene models.

### Genome Assembly and Functional Annotation of the Reference Sample

The genome of the reference *C. pini* sample was assembled from PacBio HiFi reads using hifiasm v0.16.1 (Cheng et al. 2021; Cheng et al. 2022) with default parameters. Mitochondrial and Ribosomal contigs were removed from the assembly prior quality assessment. Genome completeness was evaluated using BUSCO v5.2.2 (Manni et al. 2021), with the basidiomycota_odb10 lineage dataset. The assembly was not phased, and haplotypes were collapsed during the assembly process. Assembly statistics, including total length, N50, and contig count, were calculated using QUAST v5.2.0 (Gurevich et al. 2013). The final assembly was used for gene prediction and downstream analyses. Telomeric repeats were identified using a custom Python script that searched for at least five consecutive repeat units of either the CCCTAA or TTAGGG motif. Repeats were considered telomeric if they were ≥60 bp in length and located within 100 bp of a scaffold terminus.

Transposable elements (TEs) and other repeats were annotated using EarlGrey (Baril et al. 2024) with the option -r 2759 (Eukaryotes). Structural gene annotation was carried out using MAKER3 v3.01.03, integrating three ab initio predictors: Augustus v3.4.0, SNAP (2019-06-03) and GenmarkES v3.62. RNA-Seq data and protein homology evidence were included in all MAKER3 runs. Reference protein sets were obtained from *Cronartium quercuum* f. sp*. fusiforme* G11 v1.0 (Pendleton et al. 2014), *Melampsora larici-populina* v2.0 (Persoons et al. 2021), and *Puccinia graminis* f. sp. *tritici* v2.0 (Duplessis et al. 2011; Cuomo et al. 2017). Augustus and SNAP were iteratively trained on single-copy BUSCO genes. MAKER3 was run separately with each predictor, incorporating all evidence, and the resulting models were used for re-training. GeneMark-ES was self-trained using its default approach. A final MAKER3 run combined all three predictors with RNA-Seq and homology evidence to generate a final gene model set. Functional annotations were assigned using InterProScan v5.43 with database release 83.0. Gene models were provided with tentative descriptions based on homology to UniProt entries.

### Draft Genome Assembly of 18 *C. pini* samples

Paired-end Illumina reads from 18 *C. pini* samples were assembled de novo using SPAdes v3.15.5. Reads from two flow cell lanes per sample were quality-checked with FastQC v0.11.9 and trimmed for adapter sequences using Trimmomatic v0.36. Cleaned reads were provided to SPAdes as two paired-end libraries, using the --isolate mode and default k-mer sizes.

### Genome Feature Visualization

Genome feature plots were generated using Blobtools v4.1.5 (Challis et al. 2020) and R v4.3.2 (R Core Team 2021). The R packages tidyverse (Wickham et al. 2019), ggplot2 (Wickham 2016), and circlize (Gu et al. 2014) were used to visualize genome assembly statistics. Repeat and gene annotations were processed with GenomicRanges (Lawrence et al. 2013) and rtracklayer (Lawrence et al. 2009). Custom functions were developed to calculate window-based coverage (10 and 50 kb) of annotated genes and repeat elements. These data were visualized using circular plots with circlize and linear chromosome plots with karyoploteR (Gel and Serra 2017). Telomeric repeat counts were overlaid on scaffold features to highlight telomere localization.

### Identification and Phylogenetic Analysis of Mating-Type (MAT) Genes

Mating-type loci were identified across all *C. pini* genome assemblies using previously characterized protein sequences of HD transcription factors, STE3.2 pheromone receptors, and MFA pheromone precursors (Cuomo et al. 2017; Luo et al. 2024). These sequences were used as queries in TBLASTN searches against each genome assembly to detect homologous regions corresponding to MAT loci (Supplementary Table S3). Predicted MAT gene regions were extracted, and their coding sequences (CDS) were refined using ExPASy Translate tools and GeneWise to account for potential introns and frameshifts. Alignments were performed in MEGA to confirm accurate CDS boundaries and verify homology across samples. Additional MAT gene sequences from other rust fungi were included as reference taxa for phylogenetic inference (Supplementary Table S3). Protein sequences were used to construct phylogenetic trees with IQ-TREE v2.1.3 (Minh et al. 2020). The best-fit substitution model for each MAT gene was selected using ModelFinder, based on the Bayesian Information Criterion (BIC) (Kalyaanamoorthy et al. 2017). The optimal models identified were: JTT+F+G4 for STE3.2, bW-HD1, and bE-HD2, and JTTDCMut+G4 for MFA. Tree support was evaluated using ultrafast bootstrap approximation (-bb 1000) and approximate likelihood ratio test (-alrt 1000) (Hoang et al. 2018). Final trees were visualized using FigTree v1.4.3 (http://tree.bio.ed.ac.uk/software/figtree/) and graphically edited in Inkscape (https://inkscape.org).

### Genome-wide Coverage Analysis

PacBio HiFi reads were aligned to the reference genome using minimap2 v2.1 with the map-hifi preset. Coverage depth was calculated using Samtools depth v1.16, and coverage distributions were visualized in R v4.3.2 using ggplot2. Genome-wide coverage distributions for draft genome assemblies of the remaining 18 samples were visualized using a custom R script in R v4.3.2. Sequence information was extracted from SPAdes-generated scaffold headers in FASTA files. Coverage values were parsed and filtered to exclude small scaffolds (<200 bp) and extreme outliers (<8× or >200× coverage). For each sample, coverage histograms were generated using ggplot2, with the peak coverage bin annotated by a vertical dashed line to indicate modal genome-wide coverage. Composite panels were assembled using gridExtra to allow side-by-side comparisons. Final plots were exported as SVG files for high-resolution visualization and further editing in Inkscape (https://inkscape.org).

## Data and resource availability

The data for this study has been deposited in the European Nucleotide Archive (ENA) at EMBL-EBI under the project accession number PRJEB89018. All custom scripts used in this study are publicly available at https://github.com/paula-gomez-zapata/Mating-type-genes-in-Cronartium-pini

## Acknowledgements

We would like to thank the members of the Mykopat and Theoretical Biology & Bioinformatics (TBB) groups for their valuable suggestions and feedback, which greatly improved the quality of this manuscript. Special thanks to Katarina Ihrmark for her high-quality DNA extractions, which were essential to this work. We also extend our gratitude to Duhen, A.P.M. (Anna) for her assistance in detecting the MAT genes in the reference genome. Sequencing was performed by the SNP&SEQ Technology Platform in Uppsala. The facility is part of the National Genomics Infrastructure (NGI) Sweden and Science for Life Laboratory. The SNP&SEQ Platform is also supported by the Swedish Research Council and the Knut and Alice Wallenberg Foundation. The genome sequencing was funded by a grant to Prof. Stenlid from SLU Forest Damage Center. Dr. Paula A. Gómez-Zapata and Dr. Åke Olson’s work is funded by Carl Trygger Foundation, CTS 22:2086.

## Author contributions

Å.O. and J.S. conceptualized and developed the project. Å.O. supervised the project. P.A.G.Z. performed data analysis, interpretation and wrote the draft manuscript. C.T.R. assembled the reference genome. M.B.D., M.S., and Å.O. performed data analysis and interpretation. B.S. and J.K. collected the samples. All authors contributed to editing the manuscript.

